# A switch in the development: microRNA arm usage screening in zebrafish suggests an important role of arm switching events in ontogenesis

**DOI:** 10.1101/2021.11.05.467297

**Authors:** Arthur C. Oliveira, Luiz A. Bovolenta, Lucas Figueiredo, James G. Patton, Danillo Pinhal

## Abstract

In metazoan, regulatory molecules tightly control gene expression. Among them, microRNAs (miRNAs) are key regulators of several important features, like cell proliferation, differentiation, and homeostasis. During miRNA biogenesis, the canonical strand that loads onto RISC can be switched, in a process called “arm switching.” Due to the miRNA-to-target pairing peculiarities, switching events can lead to changes on the gene-targeted repertoire, promoting the modulation of a distinct set of biological routes. To understand how these events affect cell regulation, we conducted an extensive and detailed *in silico* analysis of RNA-seq datasets from several tissues and key developmental stages of zebrafish. We identified interesting patterns of miRNA arm switching occurrence, mainly associated with the control of protein coding genes during embryonic development. Additionally, our data show that miRNA isoforms (isomiRs) play an important role in differential arm usage. Our findings provide new insights on how such events emerge and coordinate gene expression regulation, opening perspectives for novel investigations in the area.

## 1. Introduction

Gene expression of metazoans is controlled by a myriad of molecular regulators that drive cell fate. Among them, microRNAs (miRNAs) emerged as important regulators of several biological processes, from cell development, such as gastrulation, hematopoiesis, and tissue formation (Espinoza-Lewis and Wang, 2012; Ivey and Srivastava, 2015; Chen and Chen, 2019) to homeostasis like response to environmental influence and diseases (Qiu et al., 2012; Reddy, 2015; Mohammed, 2020). MiRNAs are a large class of non-coding RNAs that regulates gene expression post-transcriptionally, mainly by interacting with the 3’UTR of the mRNA targets (Lai, 2002; Dexheimer and Cochella, 2020). Thanks to their particular binding affinity to targets, miRNAs can promote an extensive spectrum of interactions, enabling a single miRNA to regulate multiple genes, while a single gene can be regulated by several miRNAs.

During miRNA biogenesis, one strand of the miRNA duplex is selected to be loaded into the RISC (RNA Induced Silence Complex), while the other is quickly degraded (O’Brien et al., 2018; Treiber et al., 2018). The strand that is selected to be part of the mature miRNA-RISC is often the same, regardless of the tissue or developmental stage of the cell that is expressing the miRNA, being named as canonical strand. However, in specific circumstances, the miRNA-RISC selects the passenger strand to be loaded instead of the canonical, in a process called arm switching (Griffiths-Jones et al., 2011). Since miRNA regulation is driven by the complementarity between them and their targets, switching the miRNA strand that is load into RISC may change its targets repertoire and thus heavily affect the biological functions and pathways controlled by that miRNA (Marco et al., 2012). Although the mechanisms that promote the arm switching events are not understood, the thermodynamic and 5’ nucleotide identity of the duplex exert key influence (Kim et al., 2020).

Several arm switching events were reported within tissues, organs, or developmental stages of living vertebrates, including *Homo sapiens, Mus musculus, Oryctolagus cuniculus, Gallus gallus*, and *Oreochromis niloticus* (Ro et al, 2007; Glazov et al., 2008; Li et al., 2011; Guo et al., 2015; Pinhal et al., 2018). Switches in arm usage were also depicted between invertebrate species, such as, *Drosophila melanogaster* and *Tribolium castaneum* (Griffiths-Jones et al., 2011). Additionally, several studies reported an association between arm switching events and diseases or cell homeostasis unbalance, like in several types of cancer (Kuo et al., 2016; Tsai et al., 2016; Chen et al., 2018; Zhang et al., 2019).

In this paper, we perform a genome-wide microRNA arm switching screening in zebrafish to understand their impacts on gene regulation of healthy organisms. Using bioinformatics, we identified consistent patterns that associate arm switching incidence with vertebrates’ development, and establish a steady correlation between arm switching events and isomiR profile.

## 2. Results

### 2.1. Overview of RNAseq data

We gathered RNAseq data from two pre-published experiments, contemplating an extensive repertoire of both adult and developmental samples from zebrafish. Adult tissue samples were collected from SRP041544 study (Vaz et al., 2015), providing data from the ovary, testes, eye, heart, and male and female brain, gut, and liver; while embryonic samples were obtained from SRP028895 (Wei et al., 2012), containing data from 256 cells (2.5 hours post-fertilization – hpf), sphere (4hpf), shield (6hpf) and 24hpf. From these experiments, we obtained about 508 million raw reads that had their quality verified and adapters trimmed. From the 498 million remaining reads, about 163 million passed through all filtering steps and were aligned against the zebrafish genome for miRNA identification.

Collapsed reads distribution revealed that ∼70% had a range of 21 to 23 nucleotides (nts). The prevalent size for most samples were 22 nts, except for the male and female brain in which the prevalent size was 23 nts. Gonadal samples also had an additional peak at 25 nts, typical of piwi-RNAs (piRNAs). PiRNAs are abundant in gonadal tissues, acting in silencing transposons and retrotransposons (Lakshimi and Agrawal, 2008), contributing to the protection and maintenance of the integrity and stability of the gamma cell genome (Sharp, 2009).

### 2.2. MiRNA identification and detection of arm switching events

We identified 1288 mature 5p- and 3p-miRNAs expressed, originated from 644 precursors, being 578 known pre-miRNAs and 66 putative novel pre-miRNAs (Table S2 and S3).

Through a careful examination of the miRNA expression profiles, we identified arm switching events in 14 pre-miRNAs, being 13 from known pre-miRNAs (i.e., dre-mir-27b-1, dre-mir-27b-2, dre-mir-92a-1, dre-mir-92a-2, dre-mir-135b, dre-mir-137-1, miR-137-2, dre-mir-153a-1, dre-mir-153a-2, dre-mir-153b, dre-mir-222a, dre-mir-2188, dre-mir-31) and a single arm switching in the novel pre-miRNA dre-mir-n001 (Fig. 1). 5p/3p ratio values can be found at Table S4. The arm switching events identified were further divided into four biological scenarios detailed below.

**Figure 1.**
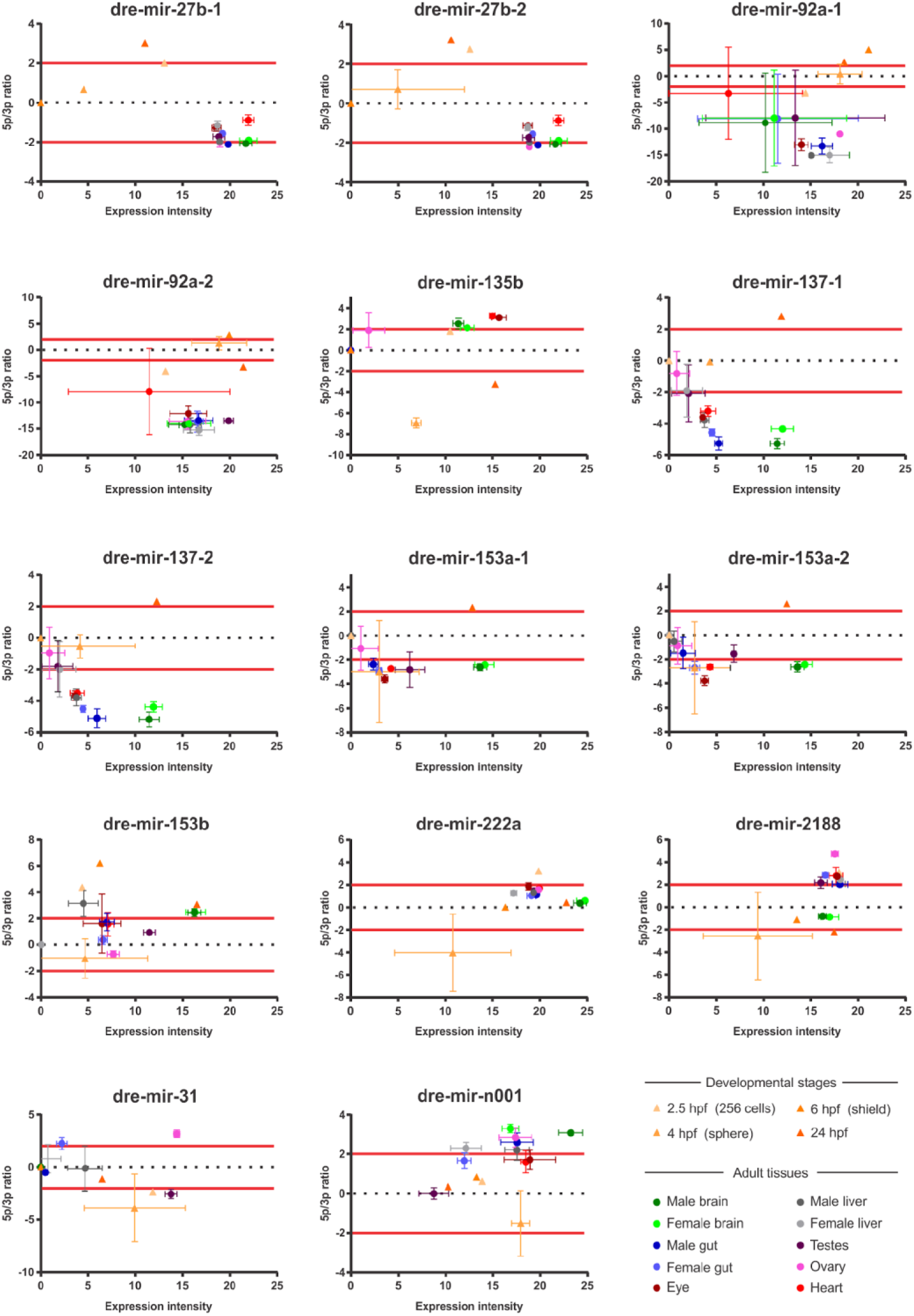
MA plot of zebrafish arm switching events. Y-axis represents 5p/3p arms ratio. Positive values represent 5p arm predominance while negative values represent 3p arm predominance. Red lines represent the 2-fold threshold for a bona fide arm switching. X-axis represents the expression intensity of the miRNAs. Triangles represent developmental stages and circles, adult tissue.

One scenario corresponds to the occurrence of events between developmental stages and adult tissues (Fig. 1). This scenario contemplates all events identified. Remarkably, half of the differential arm prevalence was between embryos 24hpf and adult brain.

In the second scenario, we identified miRNA arm switching between distinct developmental stages, as for dre-mir-27b-2, dre-mir-135b, dre-mir-192a-1, dre-mir-153a-1, dre-mir-153a-2, dre-mir-222a, and dre-mir-2188 (Fig. 2). With the exception of dre-miR-153a-1/2, in which no expression on 2.5hpf period was detected, the miRNA 5p/3p ratio of 2.5hpf period always corresponded to the ratio found on adult samples (Fig. 1). All major miRNA 5p/3p ratio changes occurred between 2.5hpf and the immediately subsequent 4hpf sphere stage. After this early developmental period, variant patterns in arm usage were observed for the distinct miRNAs.

**Figure 2.**
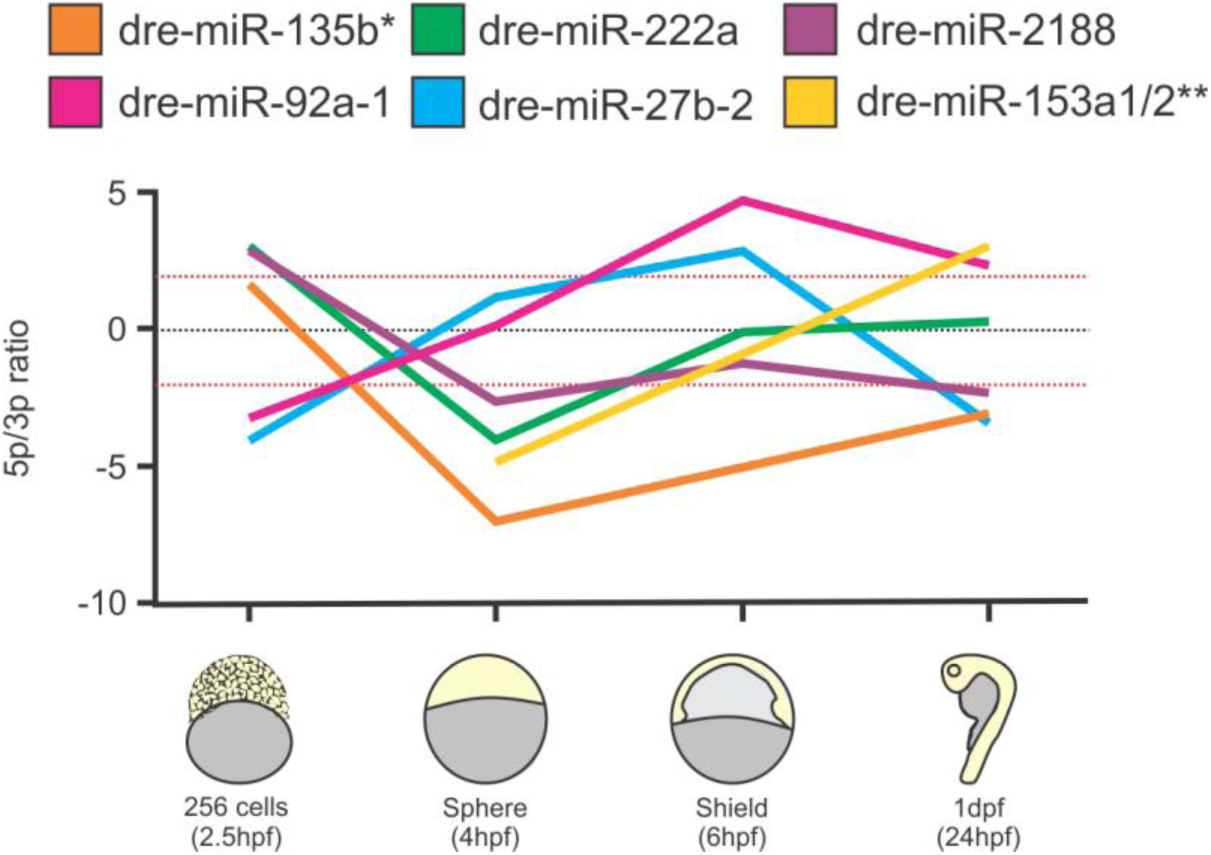
5p/3p ratio of arm switching events during the four developmental stages investigated. Red lines represent the 2-fold threshold for a bona fide arm switching.* Expression of dre-miR-135b was not detected on the 6hpf sample. ** Expressions of dre-mir-153a-1 and -2 were not detected on 2,5 and 6hpf samples.

Sex-biased variant arm usage contemplates the third scenario, with dre-mir-31 inverting its 5p/3p ratio between testes and ovaries (Fig. 1). Interestingly, this finding was a unique miRNA arm switching between two adult tissues.

The last biological scenario in which arm switching events were identified refers to two miRNA paralogs (Fig. 3). For the dre-mir-92a, there is a shift on 5p/3p ratio between dre-miR-92a-1 and dre-miR-92a-2 on 24hpf. For dre-mir-153, in brain samples, dre-miR-153a-1-3p is more expressed than its 5p counterpart, while dre-miR-153b-5p is more expressed than dre-miR-153b-3p.

**Figure 3.**
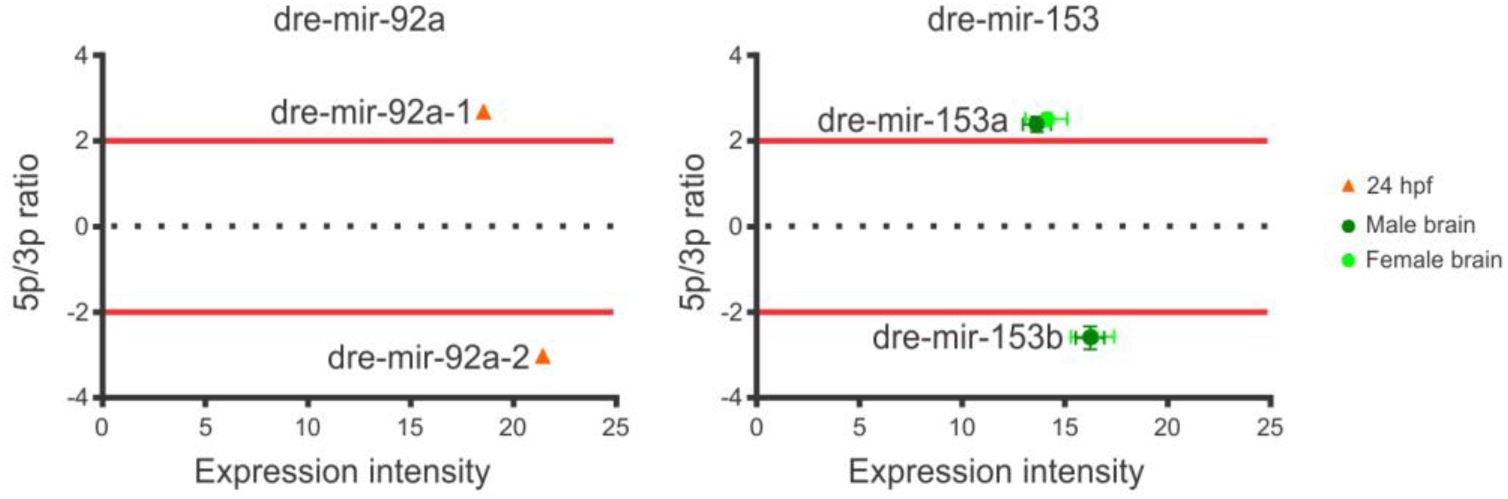
Arm-switching events between paralog copies of miRNAs dre-mir-92a and dre-mir-153. Red lines represent the 2-fold threshold for a bona fide arm switching.

### 2.3. Target prediction and functional enrichment data

Providing fifty percent of the zebrafish arm switching events were depicted between embryos 24hpf and adult brain, we exclusively performed target prediction, functional enrichment, and isomiR analysis in these tissues. This analysis allowed us to properly evaluate any prospective correlation between either enriched targets or isomiR profile with differential arm usage without external bias caused by multiple tissue deviations.

Target prediction analysis returned a total of 16,593 and 13,681 potential targets of dre-mir-27b-1/-2, dre-mir-135b, dre-mir-137-1/-2, and dre-mir-153a-1/-2 expressed in the brain and 24hpf tissues, respectively.

Similar to reports by other experiments (Griffiths-Jones et al., 2011 e Marco et al., 2012), our data demonstrated a low percentage of overlap between targets of each arm, except dre-miR-153a-1/-2, that returned a higher number of 5p arm targets than average, contemplating several shared targets with its 3p counterpart. Additionally, we did not find any correlation between the number of targets and the prevalence of the arm on a given tissue (Table 1).

**Table 1.** Representative number of predicted targets for 5p and 3p arms of miRNAs dre-mir-27b-1/-2, dre-mir-135b, dre-miR-137b-1/-2, and dre-mir-31, under distinct biological scenarios.

| miRNA | Biological scenario | 5p arm | 3p arm | Targets overlap |
| --- | --- | --- | --- | --- |
| **dre-mir-27b-1/-2** | brain | 2356 | **2465** | 1057 |
|  | 24 hpf | **1882** | 2035 | 838 |
| **dre-mir-135b** | brain | **1065** | 1461 | 378 |
|  | 24 hpf | 1302 | **1749** | 469 |
| **dre-mir-137b-1/-2** | brain | 1656 | **1358** | 448 |
|  | 24 hpf | **1298** | 1165 | 340 |
| **dre-miR-153a-1/-2** | brain | 3974 | **1733** | 1147 |
|  | 24 hpf | **3343** | 1432 | 947 |

Bold values represent the predicted targets of the most expressed arm. Target gene list these miRNAs can be found in Table S5.

Interestingly, besides the divergences in the set of miRNA targets regulated by the 5p and 3p arms, we also detected differences among functional categories targets are involved according to the prevalent arm (Fig. 4; Table S6). Indeed, the most pronounced divergences occur within molecular functions and biological pathways, while for biological processes and cellular components, although there are a significant number of processes controlled exclusively by one of the two arms, it is still possible to verify the occurrence of common terms.

**Figure 4.**
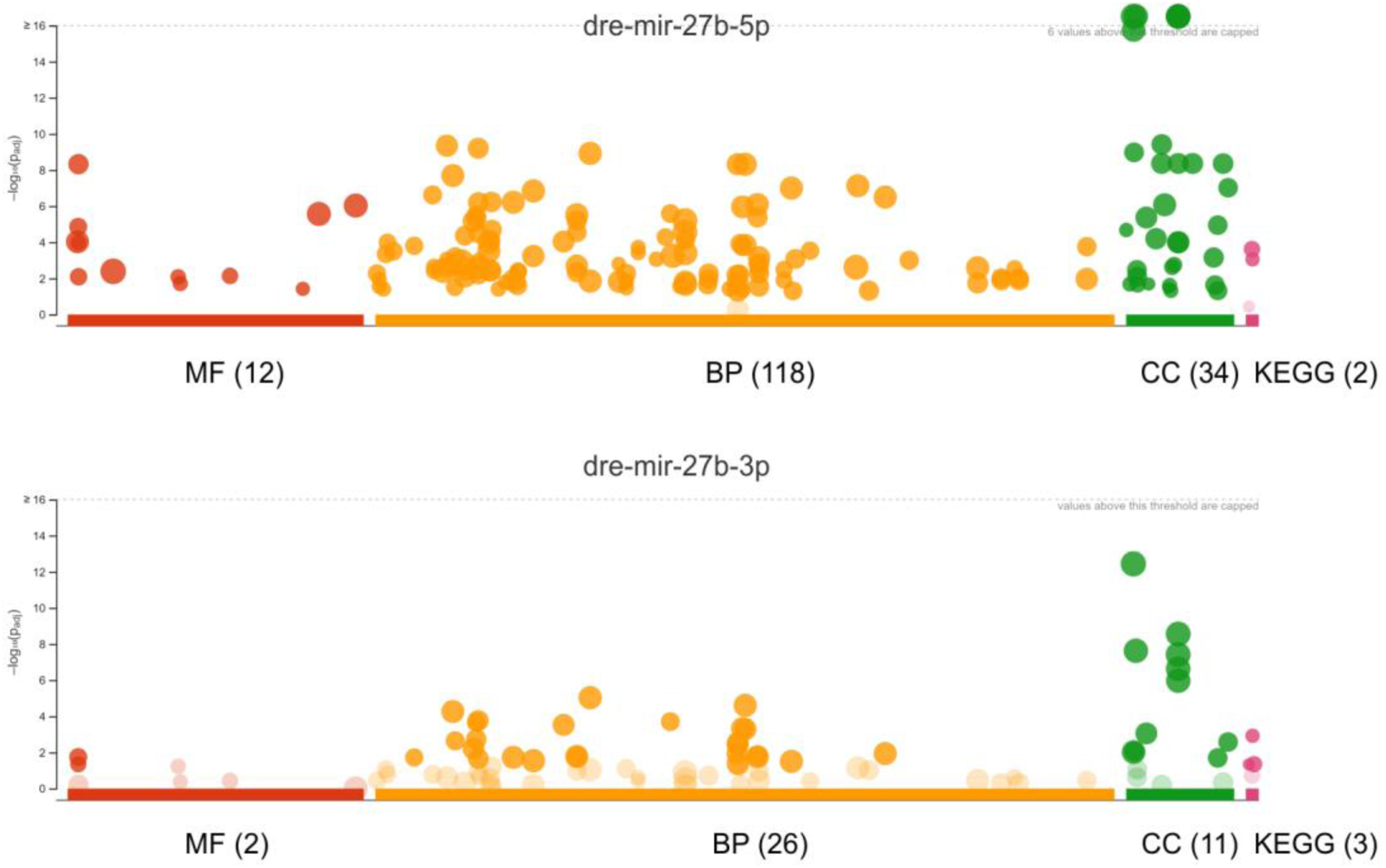
Graphic visualization of dre-miR-27b functional enrichment on embryo 24hpf. Red dots = Molecular function. Yellow dots = biological process. Green dots = Cellular components. Pink dots = Biological pathways. Point position on the Y-axis indicates P-value. Point position on the X-axis represents specific terms. Dot diameter represents the number of genes associated with the term. Light circles of each colour represent terms found without significant P-value. Detailed results from functional enrichment analyzes can be found in Table S6.

For dre-miR-27b expressed on embryo 24hpf, only one molecular function (transcription coregulator activity; GO:0003712) and one KEGG pathway term (Notch signaling pathway; KEGG:04330) was enriched on both 5p and 3p arm. Alternatively, 11 of the 34 cellular context terms enriched and 22 of the 119 biological process terms were observed on both arms. Interestingly, terms enriched only in one arm was observed only on dre-miR-27b-5p, the arm most expressed on embryo 24hpf, for molecular function, cellular context and biological process, while two KEGG terms (Insulin signaling pathway; KEGG:04910 and Terpenoid backbone biosynthesis; KEGG:00900) were enriched exclusively on 3p arm (Fig. 4; Table S6). When searching for enriched terms of dre-miR-27b expressed in the brain, where dre-miR-27b-3p is the prevalent arm, we observed four of the 21 molecular functions and one of the four KEEG (Endocytosis; KEGG:04144) terms enriched on both arms. In the cellular context and biological process, there were 17 of 46 and 37 of 115 enriched on both arms. Different from embryo 24hpf sample, there was a greater number of terms enriched exclusively to either 5p or 3p (Table S6). Results for the functional enrichment for other miRNAs on embryo 24hpf and brain tissue can be found in Table S6.

### 2.4. Correlation between arm switching events and isomiR expression

IsomiRs are variant miRNA molecules (also called miRNA isoforms) that differ from the representative miRNA by having modifications in its sequence. Such modifications include single nucleotide substitutions, insertions, deletions, 3′ end non-templated additions as well as 5′ and/or 3′ cleavage shifts (Pinhal et al., 2018). Since these variations in miRNA sequence may promote alterations on both the relative instability of the miRNA-miRNA* duplex and the identity of the initial nucleotide of each arm, we seek to analyze the isomiR patterns in the identified arm switching events to evaluate how they impact arm selection.

Our analysis identified several modifications in the sequence of expressed transcripts, both in the template and non-template read, thus characterizing typical isomiRs (Fig. 5). Among isomiRs, the great majority of isoforms probably originated from distinct Dicer and Drosha cuts during miRNA biogenesis, whereas only a small fraction of them (∼0.72%) corresponded to non-template isomiRs, such as single nucleotide modifications Interestingly, identical isomiRs were present in both embryo 24hpf and adult brain, while isomiRs exclusive to only one tissue were rare. However, we observed that the relative expression of the shared isoforms changed according to the tissue analyzed. Such variations on isomiR expression were strong enough to change the representative isoform (i.e. the isoform most expressed) of at least one of the mature arms between the tissues, occurring in five of the seven miRNAs analyzed (Table 2). These data suggest that the occurrence of arm switching is mainly followed by a shift in the representative isoform of the mature miRNAs. For example, dre-miR-135b-5p is less expressed than dre-miR-135b-3p in embryo 24hpf and its representative isoform is 23 nucleotides long. However, with the development of the brain, an arm switching event occurs in that tissue, and dre-miR-135b-5p becomes more expressed than its 3p counterpart. This event is followed by shifting the representative isoform of dre-miR-135b-5p on brain tissue, which is shorter than the one present embryo 24hpf, with a missing adenine at 3’ region (Table 2).

**Figure 5.**
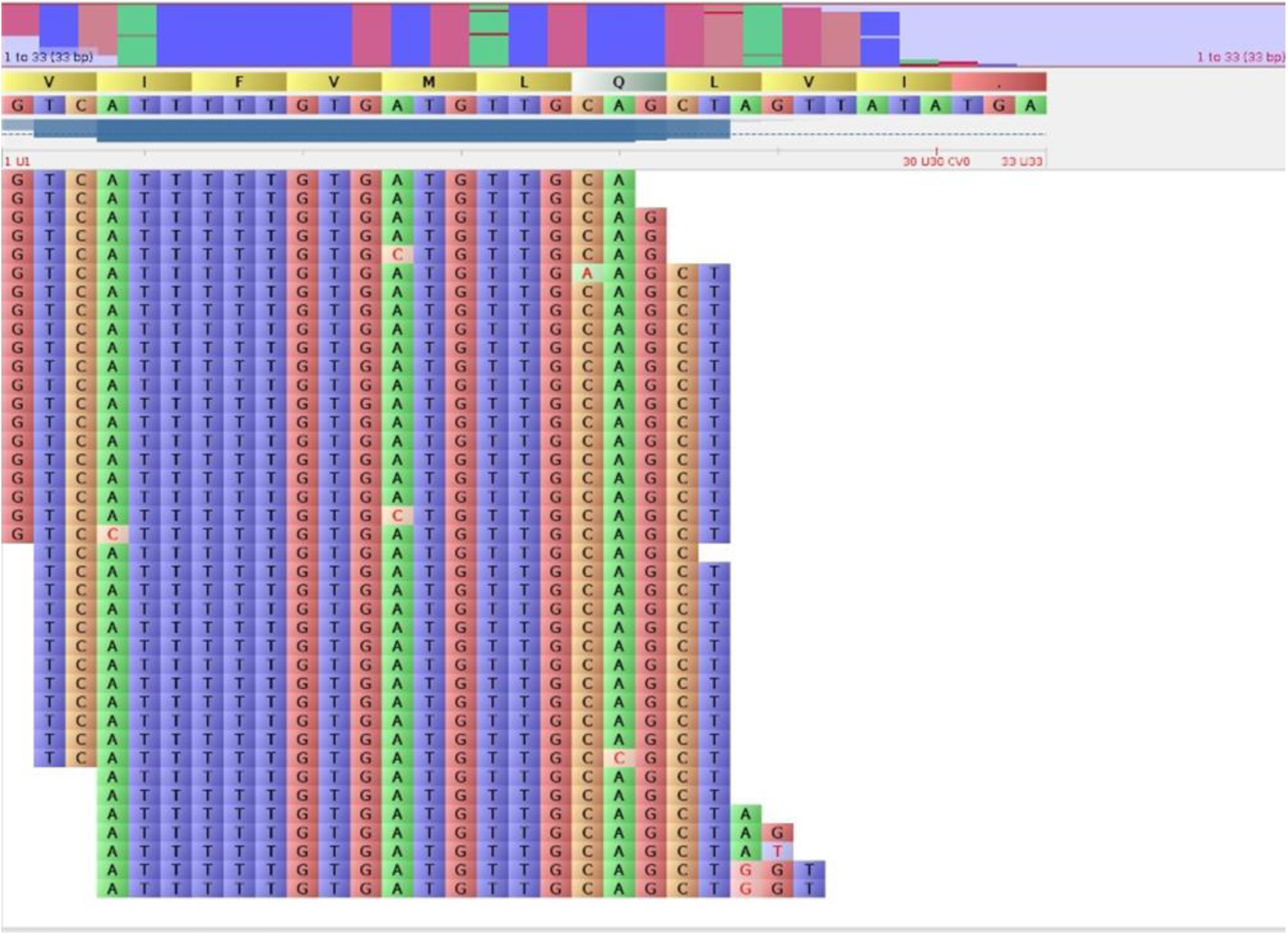
IsomiR patterns of dre-miR-153a-2 are expressed in the brain. Several patterns of isomiRs were identified on both arms, such as reads with insertions and deletions on both 5’ and 3’ ends, 3’non-templated insertions, and single nucleotides mutations.

**Table 2.**
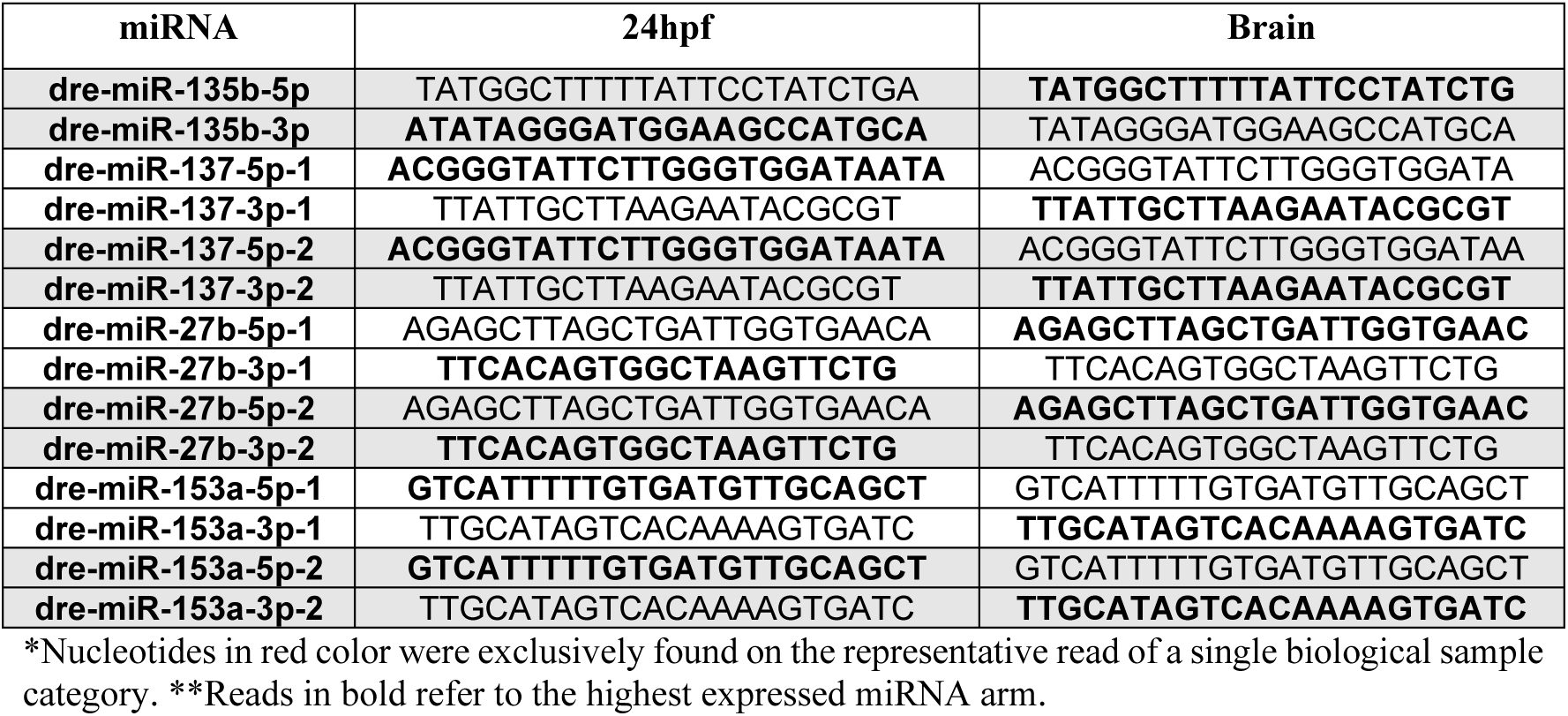
Representative isoform from 5p and 3p miRNA arms at embryos 24hpf and adult brain.

Our data also revealed specific isomiR expression patterns in each tissue. For instance, the representative isoforms of embryo 24hpf were usually longer than the representative isoforms of the adult brain. Additionally, we identified a higher occurrence of non-template isomiRs in the brain (∼1.29%) than in 24hpf (0.29%). Finally, dre-miR-135b demonstrated an interesting pattern in which the representative read of dre-miR-135b-3p has an additional adenine in its 5’ portion in 24hpf sample in relation to its counterpart in the brain sample. IsomiRs arising from variations in the 5’ region are extremely rare and promote an event called ‘seed shifting’, in which the seed region, the main responsible for target recognition, has its sequence changed (Guo an Chen et al., 2014; Haseeb et al., 2017).

## 3. Discussion

### 3.1. Arm switching is a conserved mechanism, but events are rare and mainly species-specific

Arm switching was identified in many species (Landgraf et al., 2007; Ro et al, 2007; Glazov et al., 2008; Wit et al., 2009; Chiang et al, 2010; Griffiths-Jones et al., 2011; Pinhal et al., 2018). However, in each study, a different set of miRNAs exhibited changes in arm prevalence and only a few events were identified so far. Pinhal et al. (2018) performed a large-scale analysis of miRNA on Nile tilapia (*Oreochromis niloticus*) being able to identify arm switching in 9 miRNAs, from the 368 miRNA loci identified in their analysis. The same approach was applied in the 674 miRNA loci of zebrafish and only 14 miRNA changes passed the threshold to be considered a bona fide arm switching (i.e. 2fold difference in at least two tissues). Interestingly, none of the miRNAs having changes in arm usage were common to zebrafish and Nile tilapia.

A high-throughput study of mouse miRNAs also reported 21 arm switching events (from 506 miRNAs identified) occurring between several developmental stages (ES, e7.5, e9.5, e12.5, and newborn) and testes, ovary, and brain of adult animals (Chiang et al, 2010). Again, none of the identified miRNAs was found also described on tilapia or zebrafish.

In addition to these large-scale analyses, other experiments have described arm switching in specific contexts. Landgraf et al. (2007) demonstrate that in mammals, miR-100 and miR-125 have as the most expressed the 5p arm, but the 3p arm has a dominant expression in some tissues. Glazov et al. (2008) identified four miRNAs (miR-135a-2, miR-30e, miR-219, and miR-30c) under arm switching during chicken development of days five, seven, and nine post-fertilization. Arm switching of miR-30e has also been reported to occur between stomach and spleen on mouse (Ro et al, 2007). Additionally, eight arm switching events were also identified occurring among four species of nematodes (*Caenorhabditis elegans, C. briggsae, C. remanei and Pristionchus pacificus*; Wit et al., 2009) and two between two species of insects (*Drosophila melanogaster and Tribolium castaneum*; Griffiths-Jones et al., 2011). Interestingly, Wit et al. (2009) report that while for vertebrates the overall predominant arm is the 5p, for invertebrates, such as *Drosophila melanogaster, D. pseudoobscura*, and the analyzed nematodes, the prevalent arm is usually the 3p, suggesting that a major change on arm prevalence may have occurred at some point of metazoan evolution.

An increasing number of studies have been searching for and identifying arm switching events in an increasing number of species. Despite that, it is notorious that the number of events identified comprehends only a small fraction of the miRNA repertoire described for the analyzed species. These data reveal that although arm switching is a conserved mechanism among metazoan, they have rare and occasional occurrence on a species-based level. Additionally, when searching for specific events, results show that the cases observed for one species usually cannot be identified in others (Fig. 6), demonstrating that arm switching is also, in its majority, species-specific and it may have to be related to particularities of the cells that the miRNAs are expressed, which vary widely between one species to another. An interesting exception is the miR-30 family, in which arm switching events of mir-30c were reported for chicken and Nile tilapia, while miR-30e-1 events were identified in chicken, Nile tilapia, and mouse.

**Figure 6.**
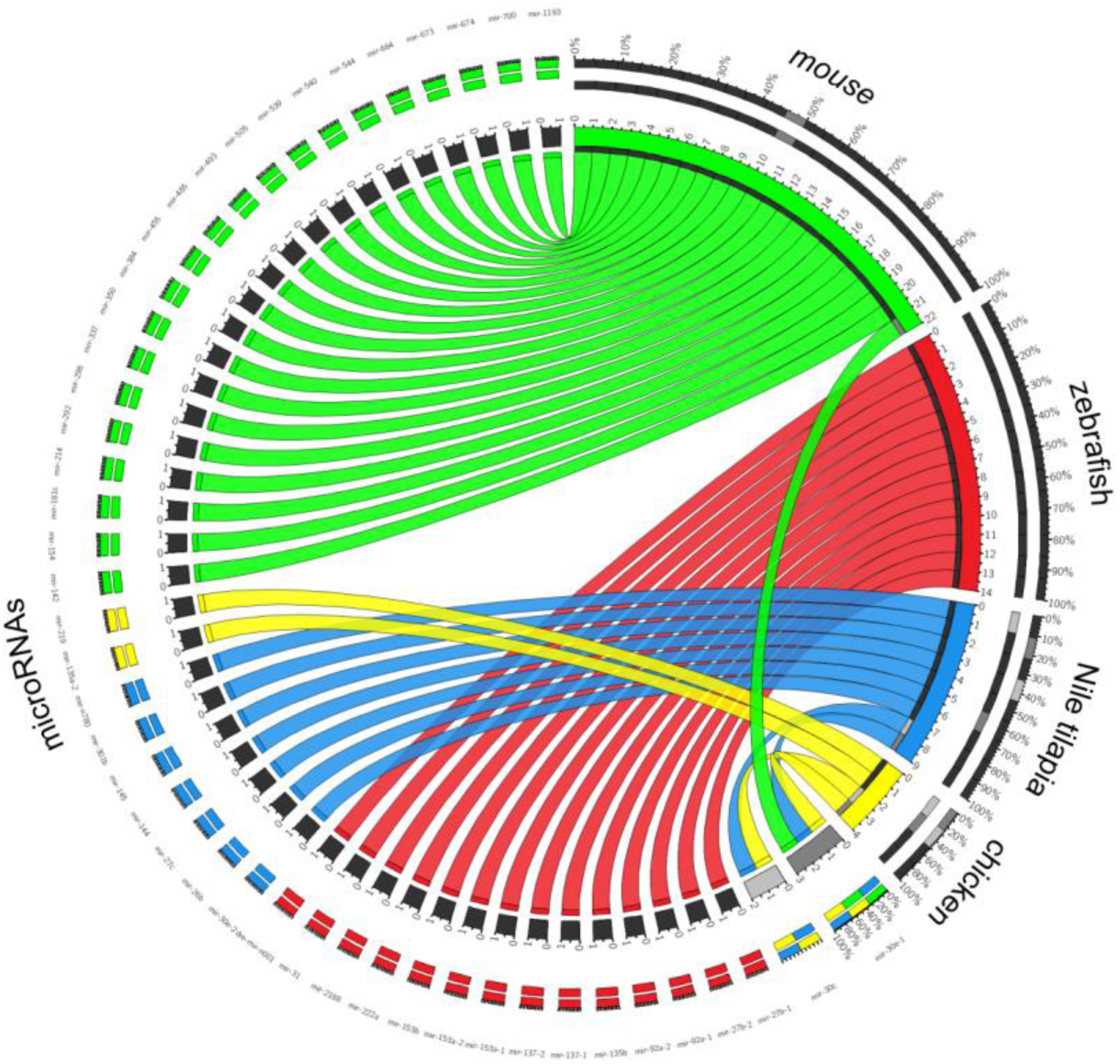
Distribution of arm switching events among species. Green, red, blue and yellow segments represent arm switching events discovered on the mouse, zebrafish, Nile tilapia, and chicken, respectively. Black squares represent arm switching events identified only on one species. Light grey square (miR-30c) represents an arm switching event identified on two species (chicken and Nile tilapia). Dark grey square (miR-30e-1) represents an arm switching event identified on two species (mouse, chicken, and Nile tilapia).

### 3.2. Arm switching is mainly related to organism development rather than tissue specification

Our analysis identified arm switching for 14 miRNAs under four biological scenarios: (i) between embryo and adult; (ii) between distinct early developmental stages; (iii) between sexes (i.e., ovary versus testis); and (iv) between miRNA paralogs, which support a major relevance of arm switching for developmental roles. Interestingly, the arm switching events identified for Nile tilapia (Pinhal et al., 2018) and mouse (Wit et al., 2009), although distinct from the ones identified here, also describe events mainly associated with the ontogenetic development of these organisms.

Strikingly, all events identified here occurred between one (or a few) developmental stages and adult tissues of zebrafish (Fig. 1). The great predominance of this class of arm switching suggests that this event can be linked mainly to the control of vital processes of the development of organisms, in which one arm acts regulating genes expressed during embryogenesis whereas, in the adult, the other arm act predominantly in genes associated with the control of tissue homeostasis. Eventually, the deregulation of this process through altered expression of the 5p and 3p arms can have harmful effects on the body.

Several studies have identified arm switching events occurring in various types of cancer (Kuo et al., 2016; Tsai et al., 2016; Chen et al., 2018; Zhang et al., 2019). In fact, epigenetic reprogramming or miRNA mutations can lead to the deregulation of several genes involved in both regulatory circuits active during development and at various stages of tumor progression, such as the Notch, Wnt, and Hedgehog pathways, (Aiello et al., 2016). Besides, several morphological changes that occurred during embryogenesis were detected in some tumors. For example, during the transition from epithelium to mesenchyme (epithelial-to-mesenchymal transition [EMT]), epithelial cells lose their characteristics, such as basal-apical polarity and cell adhesion, thus increasing their mobility, which is an extremely important event during gastrulation. The same mechanism occurs in tumor cells, increasing their mobility and facilitating the spread of cancer (Aiello et al., 2016). Future work investigating the global expression of miRNAs in embryonic cell and tumor lines can assist in confirming the correlation of the occurrence of arm switching events in both scenarios.

In addition to the discovery of arm switching between embryos and adults, our analysis detected seven events by comparing early developmental stages (Fig. 2). The developmental stages sequenced by Wei et al. (2012) and revisited by us, stand for key periods during embryonic development of vertebrates exhibiting notable variations in gene expression. In the 256-cell stage, most RNAs are from maternal origin, in which they are replaced by endogenously transcribed RNAs during the subsequent sphere stage. Ahead, during the shield stage, the formation of the three germ layers occurs, whereas at 24hpf most organs are formed.

With the exception of the paralog copies of dre-mir-153, in which no expression levels were detected at 2.5 hpf, the 5p/3p ratio at 2.5 hpf of the miRNAs always followed the ratio identified on adult tissues. Up to the period of 2.5 hpf (256 cells), the set of mature miRNAs present in the embryo is entirely made of maternal origin (Wei et al., 2012), so it is expected that the 5p/3p ratio found in this period will follow that observed in the ovary. Conversely, dre-mir-31, the miRNA in which an arm switching event between the testis and ovary was detected, is an exception, exhibiting 5p/3p ratio at 2.5 hpf opposite to that found in the ovary (Fig. 1).

In the 4hpf (sphere) stage, the embryonic machinery begins producing endogenous miRNAs, while maternal RNA starts to be degraded (Giraldez et al., 2005; Wei et al., 2012). Unsurprisingly, we found that in the 4 hpf period there is a change in the prevalence of the 5p/3p arm ratios in relation to the previous period (2.5 hpf; Fig. 3), resulting in arm switching events caused by the beginning of endogenous embryonic transcription. From that period, different patterns of arm selection were observed.

For the dre-miR-27b, for example, there was a predominance of the 3p arm over the 5p at 2.5hpf, with this ratio changing at 4hpf. However, at 24hpf, a new arm switching event occurred, in which the 3p returned to be the predominant arm. For the dre-miR-2188 and dre-miR-135b, after the occurrence of the arm switching between the periods of 2.5 and 4hpf, the 5p/3p ratio stabilized, with the prevalence of the 3p arm in the subsequent periods. Regarding dre-miR-222a, the 5p arm prevailed at 2.5hpf, but an arm switching event made the 3p arm to become the main functional transcript at 4hpf. Afterwards, from 6hpf on, both arms were equivalently expressed. For the dre-miR-92a, the complete arm switching event takes a little longer to occur. At 2.5hpf, there was a predominance of the 3p arm when, at 4 hpf period, both arms had a similar relative expression, and finally at 6 hpf the 5p arm prevailed.

An interesting situation was the identification of the arm switching of dre-mir-31, occurring between zebrafish ovary and testis (Fig. 1). Previous studies have shown that miR-31 plays an important role in differentiating gonads during embryonic development in chickens (Cutting et al., 2012). During the onset of chicken sexual dimorphism, mir-31 is more expressed in the male whereas in later stages, post-differentiation, there is a balance in the overall expression of this miRNA between the two sexes (Cutting et al., 2012). Interestingly, high levels of expression of dre-miR-31 were detected only during the early stages of development (2.5 hpf and 4hpf) and in the adult testis and ovary tissues. In stages 2.5 hpf, 4hpf and testis there are a prevalence of the 3p arm, while in the ovary, there is a greater expression of the 5p arm (Fig. 1). Although zebrafish sexual differentiation is a polygenic event, sexual reversal analyzes and DNA site mapping experiments suggest the presence of a ZZ/ZW system, similar to that found in birds (Cutting et al., 2012). Computational analysis of target prediction shows that, in birds, mir-31 regulates components of the TGF-β signaling pathway (Cutting et al., 2012), which has a fundamental role in gonadal development (Drummond, 2005; Fan et al., 2011). In this way, dre-miR-31 is likely to act similarly during the development of zebrafish sexual reproductive structures.

Finally, the last biological scenario identified in our analysis comprehends the arm switching events between miRNA paralogs. For the dre-mir-92a, there was variation in arm selection between dre-mir-92a-1 and dre-mir-92a-2 in the 24hpf. For the dre-mir-153, there was an exchange in the prevalence of the 5p and 3p arms between dre-mir-153a and dre-mir-153b in brain. In dre-mir-153a-1, the 3p arm was the most expressed, while in the dre-mir-153b the 5p arm has prevailed (Fig. 3).

The regulation of gene expression provided by miRNAs is highly conserved among organisms, but its imbalance can lead to disturbance in homeostasis. In this way, the appearance of parallel copies of a miRNA can reduce the selective pressure of maintaining the expression of a copy, facilitating the occurrence of arm switching events (Griffiths-Jones et al., 2011). According to Griffiths-Jones et al. (2011), there can be two types of arm-switching events preceded by gene duplication. The first consists of cases in which both arms are expressed. The appearance of a parallel copy can lead to the sub-functionalization of the copies, in which each copy would specialize in the regulation provided by an arm, increasing the expression of opposite arms in each copy. The second type occurs when the expression of one arm is predominant over the other. In these circumstances, gene duplication followed by an arm switching event can lead to the neofunctionalization of this new copy (Ruby et al., 2007). In both conditions, the selective pressures imposed on each copy can result in different miRNA genes, with low sequence identity, despite the common ancestry (Ruby et al., 2007). In fact, subfunctionalization and neofuctionalization derived from gene duplication events, are one of the main sources for the arising of novel miRNAs (Berezikov, 2011).

### 3.3. Targets repertoire and function changes according to the prevalent arm strand

The sequence of the 5p and 3p arms of the miRNAs differ from each other, mainly changing the seed sequence, the principal region guiding the miRNA pairing with its target mRNA (Bartel, 2009). In this sense, we proposed two possibilities in which the miRNA arms could interact with their targets: (i) each arm would regulate a different set of genes, or (ii) the arms would interact with similar genes, however at different pairing sites. Our results demonstrate a low overlap in relation to the number of specific targets, considering here the predicted targets of each arm (Table 1). These data suggest that the 5p and 3p arms of the same miRNA are mainly associated with the regulation of different biological functions. Such findings corroborate the results obtained by other researchers (Griffiths-Jones et al., 2011 and Marco et al., 2012), who also detected a low overlap in the target list of the analyzed miRNAs.

Griffiths-Jones et al. (2011) highlight that for the miRNAs evaluated in their work (miR-10, miR-993, miR-100, and miR-125), they found fewer predicted targets for the canonical arm than for the other less expressed arm and argues that perhaps there is selective pressure against the presence of matching sites for the most expressed arm. However, our data show that such pressure is not true in all situations. The dre-miR-27b-3p (brain), dre-miR-135b-3p (24 hpf), dre-miR-137b-5p (24 hpf), and dre-miR-153a-5p (24 hpf) arms have more predicted targets than their other counterpart, although they are the most expressed in these tissues. Conversely, the dre-miR-27b-5p (24hpf), dre-miR-135b-5p (brain), dre-miR-137b-3p (brain), and dre-miR-153a-3p (brain) arms are canonical and have the least number of targets (Table 1).

The divergences in the results obtained between the two studies can be associated with the methodology used in target prediction. In their work, Griffiths-Jones et al. (2011) considered the global prediction of these miRNAs, without considering their presence/absence in specific tissues, whereas our data compare tissues with each other and thus show that this ratio between 5p and 3p can vary depending on the tissue in which the miRNA is expressed.

For the miRNAs described in Table 1, in one tissue, the canonical arm has fewer targets than the least expressed arm (dre-mir-27b – 24 hpf; dre-mir-135b – brain; dre-mir-137b – brain; dre-miR-135a – brain), however, in the other tissue, the canonical arm has a greater number of predicted targets than the less expressed arm (dre-mir-27b – brain; dre-mir-135b – 24 hpf; dre-mir-137b – 24 hpf; dre-miR-135a – 24 hpf). These data demonstrate that the number of targets/miRNA expression depends on the cellular context in which it is inserted and there is not necessarily a selective pressure against canonical arm interaction sites in all scenarios.

When coming to the biological roles derived from the targets of each arm, our analysis showed that the biological processes, pathways, and molecular functions in which these targets are involved also differ according to the prevalent arm. Additionally, despite the higher similarities found in biological process terms, more pronounced divergences occur in the categories molecular functions and biological pathways. These results demonstrate a prevalent regulatory separation exercised by the 5p and 3p arms (despite of similarities between some of the biological processes regulated by both arms), because the molecular functions regulated, as well as the various correlated biological pathways, tend to be regulated by distinct arms.

It is important to highlight that the occurrence of highly enriched terms in both arms (eg developmental process (GO: 0032502) and anatomical structure development (GO: 0048856) present in the miRNA enrichment data dre-miR-27b - 24 hpf) can derive from a recurrent methodological bias of the functional enrichment technique. In the Gene Ontology consortium, each gene can have several associated terms, in which each of these terms has one or more parental terms associated with it. Therefore, the enrichment of distinct specific terms can lead to the occurrence of enrichment of parental terms in common in both arms.

Another interesting pattern detected is that, despite the differences found in the molecular functions and biological processes of the targets, the cellular components in which the targets of both arms act tend to have a greater degree of similarity (Fig. 4). This characteristic agrees with the biogenesis process of a miRNA, in which both arms derives from the same pre-miRNA, strengthening the hypothesis that the occurrence of arm-switching is due to the changes that occurred during the biogenesis of the miRNA.

### 3.4. IsomiRs are related to appearance arm switching events

The observation that variations in the 5p/3p ratio among tissues are usually linked to changes in the representative isoform of at least one arm suggests that isomiRs impact on arm switching formation. Modifications on representative isoform signature can promote the modulation of both the relative instability of the ends of the duplex and the identity of the initial nucleotide of each arm, which is known to be key features of arm selection by RISC (Kim et al., 2020).

In fact, several studies have established that changes during distinct points of miRNA biogenesis can lead to the production of duplexes with variable instability and, ultimately, promote variations in arm usage. Wu et al. (2009) demonstrated variations as early as DROSHA alternative processing on pri-miRNAs is sufficient to produce mature miRNAs with variable end instability and such variations are sufficient to promote changes in arm usage. Another study reported that the basal and apical junctions of pri-miRNAs are important to define DROSHA cut, and modifications on these regions can promote alternative DROSHA cleavage sites (Ma et al., 2013).

In a recent report, a study reported that an arm switching event of miR-324 was indeed regulated by alternative processing during its biogenesis (Kim et al., 2020). A uridylation of the pre-miRNA by terminal uridylyl transferases TUT4 and TUT 7 shifts de position of the DICER on the pre-miRNA, changing the cleavage sites (Kim et al., 2020). Such alternative processing produced non-template isomiRs, generating miRNA-miRNA* duplex in which the 3p arm is selected instead of its 5p counterpart.

Together, the data reported by these authors and our findings regarding the correlation between isoform prevalence and arm usage variations demonstrate an important role of isomiRs in promoting arm switching.

## 4. Conclusions

Our analysis demonstrated that arm switching is a conserved mechanism that is mainly assigned with vertebrate’s development. However, their particular cases tend to occur in a species- and context-specific manner. The fast modulation on the expression rate of the 5p and 3p arms and their potential to regulate different sets of biological functions, depending on the most expressed arm, enhance the control and maintenance of the cell promoted by miRNAs during the intense variation in the occurring gene expression during the development of organisms.

The differential expression of isomiRs and the alteration of the canonical miRNA isoform in different tissues is an important factor in the occurrence of these events. Additionally, this differential expression of isomiRs presents itself as an enhancer of the regulatory role of arm switching events, expanding the range of biological processes and pathways controlled by miRNAs.

## 5. Material and Methods

### 5.1. RNAseq data collection

To identify the arm switching profiles, pre-published RNAseq data were obtained from several adult tissues and developmental stages of zebrafish. Adult tissue samples were collected from SRP041544 study (Vaz et al., 2015), providing data from the ovary, testes, eye, and heart, and male and female brain, gut, and liver (n=3). Embryonic samples were obtained from SRP028895 (Wei et al., 2012), which contains data from 256 cells (2.5 hours post-fertilization – hpf) (n=1), sphere (4hpf) (n=2), shield (6hpf) and 24hpf (n=1).

### 5.2. Sample treatment, miRNA identification and 5p/3p arms characterization

Firstly, we run a quality control analysis to ensure data quality using the FastQC tool (v0.11.5; Wingett e Andrews, 2018). After, adaptor sequences were trimmed using cutadapt tool (v1.14; Martin, 2011), and the reads were converted from fastq to fasta using FASTX-Toolkit (v2.8.1). Reads were then counted, collapsed to unique reads, filtered by length (16 to 23 nts) and complexity (≥3 distinct nucleotides), and reads matching other non-coding RNAs, such as rRNAs and tRNAs were removed, using the filter module of UEA small RNA Workbench (v3.2, release 19; Stocks et al., 2012).

Treated sequences were mapped to zebrafish genome vGRCZ10 using Mapper script (v2.0.0.7; An et al., 2013), and miRNAs were identified with miRDeep (v2.0.0.7; An et al., 2013). To perform miRNA identification, pre- and mature miRNAs from zebrafish and mature miRNAs from other species were obtained from miRBase (v22; Kozomara and Griffiths-Jones, 2011) and MirGeneDB (v2.0; Fromm et al., 2020) and used as a reference, allowing a maximum of 1 mismatch outside the seed region.

Once miRNAs were identified, 5p and 3p arms from each miRNA were characterized. In this approach, pre-miRNAs were divided into two sub-sections with 30 nucleotides overlap. Each identified read were then mapped in one of these two sections using the Shortstack tool (v1.1.2; Axtell, 2013), and classified as 5p or 3p arm according. Reads mapped only to the overlap sections were filtered and removed from the analysis.

### 5.3. Data normalization and arm switching events identification

The expression of the identified 5p and 3p reads from all samples were normalized with the TMM method (trimmed mean of M values; Robinson and Oshlack, 2010) using edgeR package (R Bioconductor). After normalization 5p and 3p expression ratios of each miRNA were obtained and arm switching events were identified.

For the arm switching identification, we only considered events in which we observed a 2-fold difference between 5p and 3p arms in at least two tissues, changing the predominant arm between them and having at least 10 mapped reads on the raw read counts on both tissues (Pinhal et al., 2018).

### 5.4. Target prediction and functional enrichment analysis

The target prediction analysis was only applied to miRNAs in which arm switching events were identified. In this analysis we used the combined results from miRanda (Enright et al., 2003; www.microrna.org), RNA22 (v2, Miranda et al., 2006; https://cm.jefferson.edu/rna22/) and TargetsScan (Agarwal et al., 2015; www.targetscan.org), ensuring the best performance regarding specificity and sensibility of the analysis (Oliveira et al., 2017). The results were further filtered by the presence of the predicted targets in the tissue of interest. To do this, RNAseq data from brain and 24hpf samples were obtained and the FPKM (Fragments Per Kilobase Million) of target mRNAs were calculated. We considered only predicted targets of FPKM ≥ 5 (Soh et al., 2015; Staton et al., 2017), and mRNAs with vales lower than that were discarded.

Functional enrichment analysis of the predicted targets was performed using g:Profiler tool (Reimand et al., 2016) using the multi-query function and the significance threshold method g:SCS threshold (0.05) searching for Biological Process (BP), Cellular Context (CC), and Molecular Function (MF) (Gene Ontology; Harris et al., 2004), and Biological Pathways (KEGG; Kanehisa et al., 2000).

### 5.5. IsomiRs identification

Bam files obtained from Shortstack alignment were loaded in Tablet software (Milne et al., 2013) for graphical visualization of the reads and identification of canonical reads and isomiR patterns. Pre-miRNAs subsections generated during miRNA 5p and 3p miRNA characterization were used as reference.

## 8. Credit author statement

Arthur Casulli de Oliveira: Conceptualization, Data curation, Formal analysis, Investigation; Methodology, Writing - original draft. Luiz Augusto Bovolenta: Data curation, Formal analysis, Methodology, Writing - review & editing. Lucas Figueiredo: Data curation, Formal analysis, Writing - review & editing. Danillo Pinhal: Conceptualization, Funding acquisition, Project administration, Resources, Supervision, Writing - review & editing.

## Supplementary Material

Supplementary tables can be found at https://drive.google.com/file/d/1tdR0FnFqJXvw68Q6XcnmY7zNdfaP_lbE/view?usp=sharing

